# Multivariate Mutual Information based Feature Selection for Predicting Histone Post-Translational Modifications in Epigenetic Datasets

**DOI:** 10.1101/2025.05.28.656539

**Authors:** V. K. Dhanasekhar, Sibi Raj B. Pillai, Nithya Ramakrishnan

**Affiliations:** Institute of Bioinformatics and Applied Biotechnology, India; Indian Institute of Technology Bombay, India – 400076

## Abstract

Mutual information (MI) has been traditionally employed in many areas including biology to identify the non-linear relationships between features. This technique is particularly useful in the biological context to identify features such as genes, histone post-translational modifications (PTMs), transcriptional factors *etc*. In this work, instead of considering the conventional pairwise MI between PTM features, we evaluate multivariate mutual information (MMI) between PTM triplets, to identify a set of outlier features. This enables us to form a small subset of PTMs that serve as principal features for the prediction of the values of any histone PTM across the epigenome. We also compare the principal MMI features with those from the traditional feature selection techniques such as PCA and Orthogonal Matching Pursuit. We predict all the remaining histone PTM intensities using XGBoost based regression on the selected features. The accuracy of this technique is demonstrated on the ChIP-seq datasets from the yeast and the human epigenomes.The results indicate that the proposed MMI based feature selection technique can serve as a useful method across various biological datasets.

## 1. INTRODUCTION

Epigenetic markers including DNA methylation, histone post-translational modifications (PTMs), non-coding RNAs *etc*. regulate the expression of genes without altering the genetic sequences [1–3]. They are inherited across cell generations in a reasonably stable manner and influenced by external factors such as diet, environment and stress, leading them to be reversible [4–8]. Histone PTMs are a set of prominent markers whose patterns across the chromatin play significant roles in gene activation or repression [9–12]. Certain histone PTMs such as H3K27me3 are known to silence the chromatin region in which they are enriched while a few others like H3K36me3 and H3K4me3 are known to activate the underlying gene for transcription [13–17]. While some of these histone PTMs are conserved from yeast to mammals, several others vary in their occupancy and functional profiles across the species [18–20]. Histone PTMs and their distributions over the genes are critical for understanding their central role in chromatin remodelling and transcriptional regulation [21–23]. The combinatorial pattern of histone PTMs, hypothesized as the histone code [24–26], is known to be crucial for several biological processes downstream.

The advancement of high-throughput sequencing technologies has generated a wealth of data, creating opportunities to explore the regulatory patterns underlying the PTMs [27–29]. The emergence of machine learning, statistical, and probabilistic techniques has shown promise in capturing the combinatorial patterns from large datasets [30–35]. Pairwise correlations among different PTMs were observed and analyzed in some of the seminal works on PTM measurements [36–38]. For unbiased Gaussian distributed data, the covariance matrix indeed characterizes the statistical interrelationship between the variables. However, the complex non-linear relationships among PTMs pose significant analytical challenges in interpretability, necessitating diverse computational techniques, particularly towards the identification of the critical PTMs involved in the histone code. For example, while it is straightforward to identify the PTM that is most correlated to a given one, finding a set of *k* PTMs that predicts the given PTM sequence with minimum average error is much more combinatorially involved. The task is further complicated by the non-Gaussianity of the underlying data. Our work here focuses on identifying a set of PTMs that can predict several other PTM sequences, up to some level of accuracy. We call such a set the independent PTM feature set and term each individual PTM in this set as an independent feature.

Notice that the independent features mentioned above has some similarities to the principal components (PC). Principal Component Analysis (PCA) is a statistical dimensionality reduction technique widely used to understand the relationship between features particularly in a high dimensional space [39, 40]. The principal components are usually obtained by eigen value decomposition of the covariance matrix. The PCA effectively captures the linear relationship between the participating variables, however each PC can be a combination of several data vectors. Our approach here differs in these aspects, as we seek an independent feature set containing some of the PTM sequences themselves, and we wish to characterize relationships which are possibly nonlinear.

In this work, we develop a novel approach to obtain an independent feature set of PTMs, which could be used to predict the intensities of any other histone PTM by harnessing measures from Information theory. We validate the above technique by using ChIP-seq datasets consisting of multiple histone PTMs from the yeast and human model organisms. [41, 42]. Additionally, a comparison with selected features from other mathematical techniques such as PCA and dictionary-based reconstruction algorithms, has been provided.

## 2. MODEL AND METHODS

### 2.1. Pairwise Mutual Information (MI) based Feature Selection

Mutual information is a powerful measure for characterizing the dependence of two random variables. The MI between the random variables *X* and *Y* is calculated using:

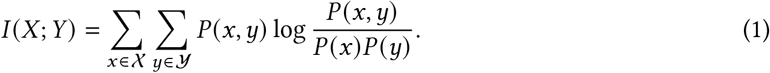

This can be contrasted with the conventional correlation measures used in PCA, which primarily targets linear relationships, by an example. Consider a zero mean Gaussian random variable *X* and an independent random variable *U* equiprobable in {−1, +1}. For *Y* = *U X*, it is easy to show that E[*XY*] = 0, thus *X* and *Y* have zero correlation. However, the mutual information *I* (*X*; *Y*) is unbounded in this case, suggesting a strong dependence. Many studies used MI-based methodologies to find insights from biological datasets. Arcos et al. [43] used a weighted MI technique to study the co-evolutionary phylogenetic relationships between influenza virus polymerase sequences - this helped the authors reduce the computational complexity in the phylogenetic inferences. Gloor et al. [44] used MI on multiple sequence alignment (MSA) to identify non-conserved sequences which are responsible for functional changes. Similarly Buslje et al. [45] used MI to identify co-evolving amino acid sequences for MSA enabling them to tackle the challenges imposed by sequence noise and redundancy normally seen phylogeny computations. Moreover, Barman and Kwon [46] created a MI-based boolean gene regulatory network (GRN) to efficiently construct regulatory networks of given target genes and to study their dynamics.

Replacing the covariance matrix in PCA by a matrix containing pairwise MI values is a potential alternative to conventional PCA, however we obtain significantly better results by resorting to the approach described in the coming section.

### 2.2. Multivariate Mutual Information (MMI)

Since the independent feature set that we seek has multiple vectors, it is imperative to move beyond the conventional pairwise relations like covariance or pairwise MI. Multivariate mutual information (MMI) is a natural candidate towards this end [47, 48]. It extends the relationship between the pairwise PTMs to multiple-tuples - in particular, we employ the three-variable MMI for our study, calculated using the following equation:

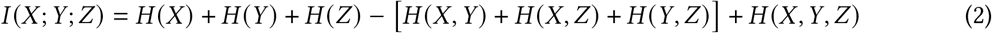

Eq. 3 can be rewritten in terms of MI as

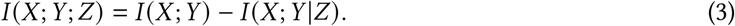

The MMI can effectively identify the graphical statistical structure of the underlying variables. To illustrate, consider two independent and identical Gaussian random variables *U* and *V*. It is easy to show that *U* +*V* and *U* − *V* are uncorrelated and independent. Consider the random vector (*X, Y, Z*) = (*U* + *V, U* − *V, V*). While *I* (*X*; *Y*) = 0 for this choice, the MMI *I* (*X*; *Y*; *Z*) is −v*e* and unbounded. This may appear to be an extreme example, and real life data is often non-Gaussian as well. Nevertheless, there are some triplets of PTMs for which MMI takes highly negative values, whereas the pairwise MI values are all positive. For example, the MMI triplet (*X, Y, Z*) = (H3K4ac, H4K16ac, H4R3me2s) has a highly negative MMI value of −0.48, whereas the corresponding pairwise MI values are all positive, as illustrated graphically in Fig 1. Based on Eq. 3, this suggests that knowing the value of *Z* makes *X* and *Y* more dependent. Thus, choosing triplets with significant negative MMI for the feature set seems to be a reasonable idea to build an independent feature set.

**Fig. 1.**
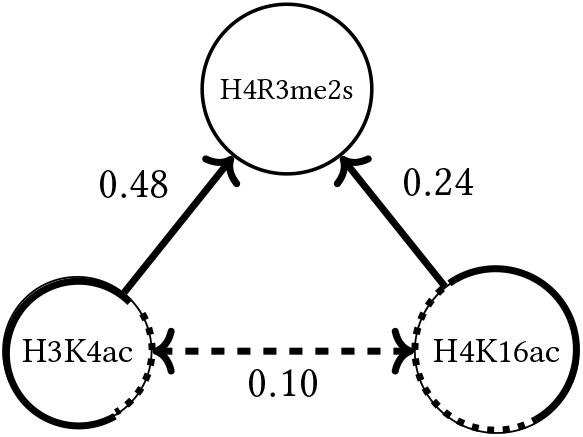
**I**llustration of a negative MMI case. Nodes in the figure represent the random variables *X, Y*, and *Z* in Equation 3, corresponding to histone modifications H3K4ac, H4K16ac, and H4R3me2s, respectively. Edge weights represent MI values.

### 2.3. Prediction of histone PTM values using XGBoost

Once a feature set is chosen, our aim is to predict the remaining PTMs using the feature set. This will bring the PTM interdependencies to the fore. To validate the features selected using the proposed techniques, we used a regression model (XGBoost, Version 2.1.3) to predict the intensities of each histone PTM (target) using the selected histone PTMs (features). XGBoost was chosen since it is well known for its efficient gradient boosting algorithms and scalability, and is particularly suited for handling complex and large datasets [49–51]. The XGBoost model parameters used were: n_estimators = 1000, max_depth = 10, random_state = 42, and n_jobs = −1.

For the prediction of target histone PTMs, the data was split into 33% (for testing) and 67% (for training the model). We employed 10-fold cross-validation using the scikit-learn Python library. To evaluate the model’s prediction accuracy, we used the coefficient of determination (*R*^2^ score) [52, 53].

### 2.4. Comparison with Linear Predictive Models

In linear models, one can think of the feature set as a matrix *A*, with each column a selected feature vector, or principal component in case of PCA. In our case, *A* is a tall matrix with more rows than columns. A target PTM sequence *y* is then approximated as *ŷ* = *Ax*, for some suitable vector *x*. For PCA, we choose a fixed number of dominant principal components to form *A*. For performance comparison, we also consider an alternative scheme in which the set of features is chosen in a greedy manner, as a function of the target PTM *y* itself. In particular, Orthogonal Matching Pursuit (OMP) is a well-known greedy technique employed for feature selection in high-dimensional datasets [54, 55]. While OMP is more suited to the cases where *A* has more columns than rows, here we employ OMP to iteratively select the most correlated features for each target until the residual error is minimal.We compared the features selected using the proposed MMI techniques against the ones obtained through OMP, for each target histone (using XGBoost regression); The selected features are presented in Fig. 3; Fig. 3(a) is a non-symmetrical heatmap of target histone PTMs and feature histone PTMs identified through OMP while Fig. 3(b) heatmap of highly negatively MMI histone PTM triplets identified using the MMI method (see Sec. 2.2 for the negative MMI case). The x-axis represents the first and second histone PTMs, and the y-axis represents the third histone PTM. The values represent the MMI scores for each triplet It is has to be noted that the OMP based features are tailored for each target histone PTM. while the MMI based features (Fig. 3(b)) are for the global set of PTMs.

Additionally, comparative analyses of the MMI-based selected features with those obtained from the traditional techniques such as PCA, Pseudo-inverse were also performed. More details about the OMP, PCA, and Pseudo-inverse pipelines are provided in the Supplementary Information (SI) Sec. 1. The overall workflow of our study is illustrated in Figure 2.

**Fig. 2.**
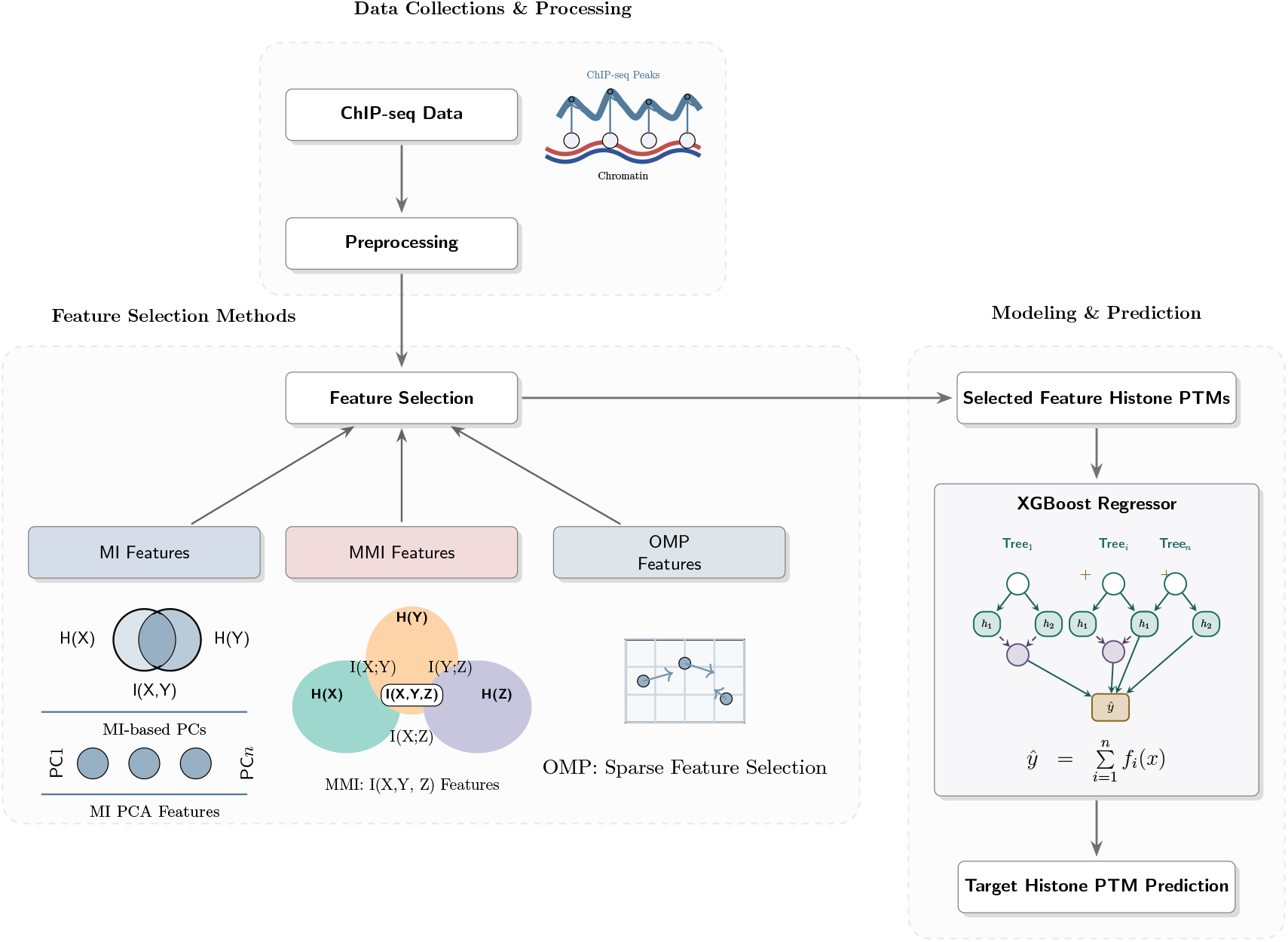
Histone PTMs analysis workflow. Preprocessed ChIP-seq data were subjected to PCA, MMI and OMP based feature selection processes. XGBoost was used to predict target histone PTMs based on feature histone PTMs selected through those feature selection methodologies.

**Fig. 3.**
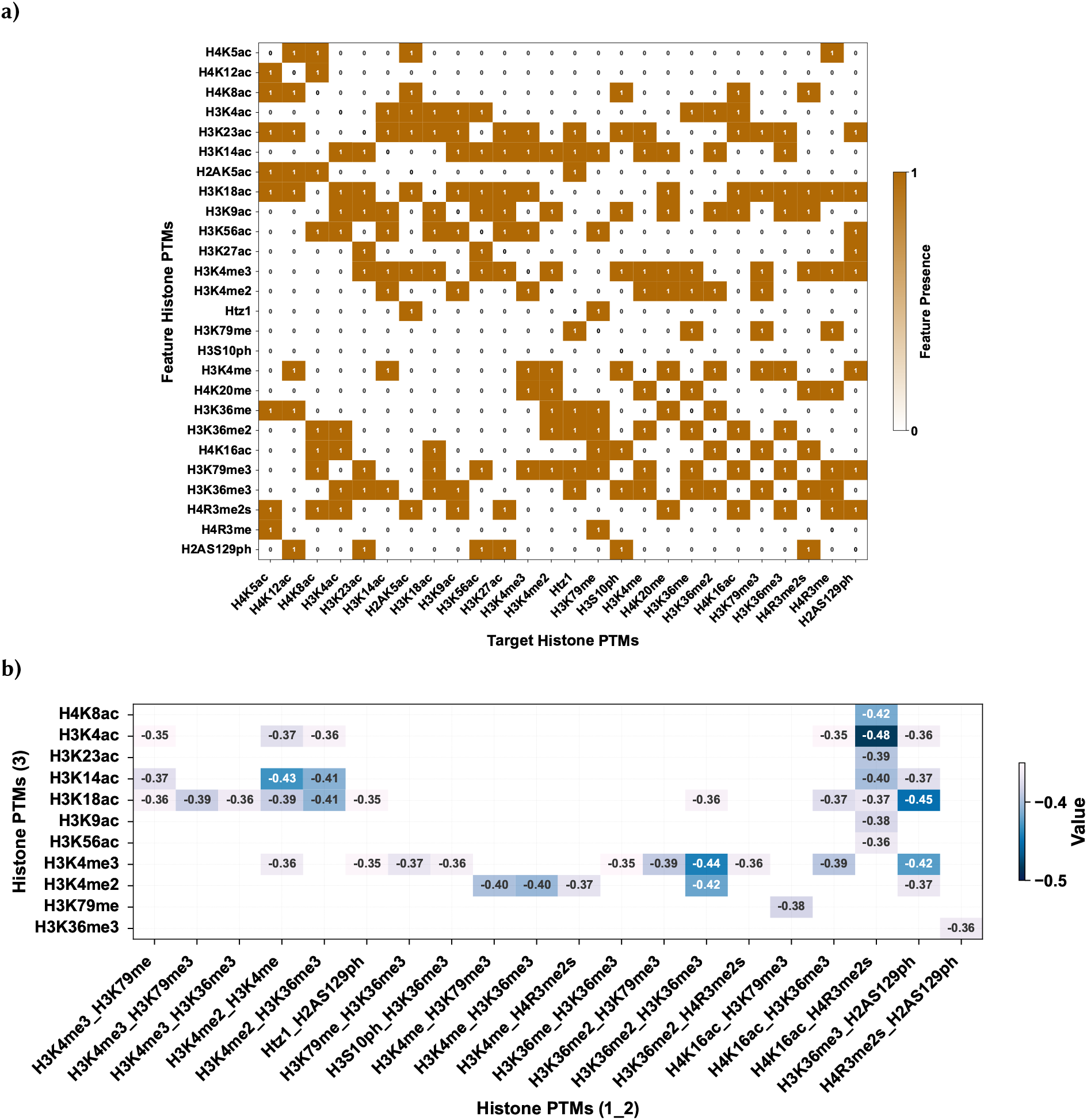
Yeast feature histone PTMs. (a) Heatmap of feature histone PTMs identified using the OMP method (see Sec. 2.4). The x-axis represents feature histone PTMs, and the y-axis represents target histone PTMs. The values (0 for absence and 1 for presence) indicate the occurrence of feature histone PTMs. (b) Heatmap of highly negatively MMI histone PTM triplets identified using the MMI method (see Sec. 2.2 for the negative MMI case). The x-axis represents the first and second histone PTMs, and the y-axis represents the third histone PTM. The values represent the MMI scores for each triplet.

## 3. RESULTS

### 3.1. Data Collection and Preprocessing

In this study, we have utilized the histone PTMs obtained from the ChIP-seq data corresponding to the histone yeast (GEO ID: GSE61888) and human (SRA ID:SRA000287) epigenomes, as reported in [41] and [42] resepectively. These datasets encompass acetylation, methylation and phosphorylation of histones and span 26 yeast histone PTMs and 39 human histone PTMs. As a preliminary step, we analyzed the nature of the distribution of datasets. Particularly, using the Shapiro-Wilk test [56, 57] the normality of the histone PTMs distribution was assessed - the results are provided in SI tables S1 and S2. It can be inferred from the test statistics and p-values that the human histone PTMs are not normally distributed while the yeast histone PTMs show significant deviations from the normality.

Next, we standardized both the datasets to zero mean; for human dataset the extreme values were trimmed - more details are provided in Sec. S3. The histograms of the histone PTMs for the two datasets before and after these steps are provided in SI figures S2-S5. Subsequently, to compute MMI, we discretized the pre-processed datasets by binning each histone PTM distribution into 100 bins with a width of 0.1 each, where the global maximum was 4 and the global minimum was −6. For the human dataset, we used 130 bins with a width of 0.56 each - here the global maximum was 40 and the global minimum was −33. We verified that the chosen bin-size is optimum by plotting the entropies for varying bin-sizes and observing minimal changes in the entropy (see SI Sec.4).

### 3.2. Results from pairwise MI and PCA

As a baseline comparison, the heatmaps and scree plots for the covariance and MI based PCA are plotted in Fig. 4 and Fig. 5 for the yeast and human datasets respectively. To assess the predictive power of the MI- and covariance-based principal components for target histone PTMs, we selected features based on the first five PCs (after transformation - for more details, see SI Sec.*S*1.2) to compute the R^2^ scores for all the histone PTMs. Figs. SI S5 (a) and (b) present the prediction results for both yeast and human datasets, respectively. Both the Figs. SI S5 (a) and (b) compare these PC-based predictions with two alternative R^2^ scores: one derived from the most highly correlated histone PTM, and another, using all remaining histone PTMS. From these analyses, it is evident that the PC-based R^2^ values are lower than those obtained using the highest-correlated histone PTM-based method.

**Fig. 4.**
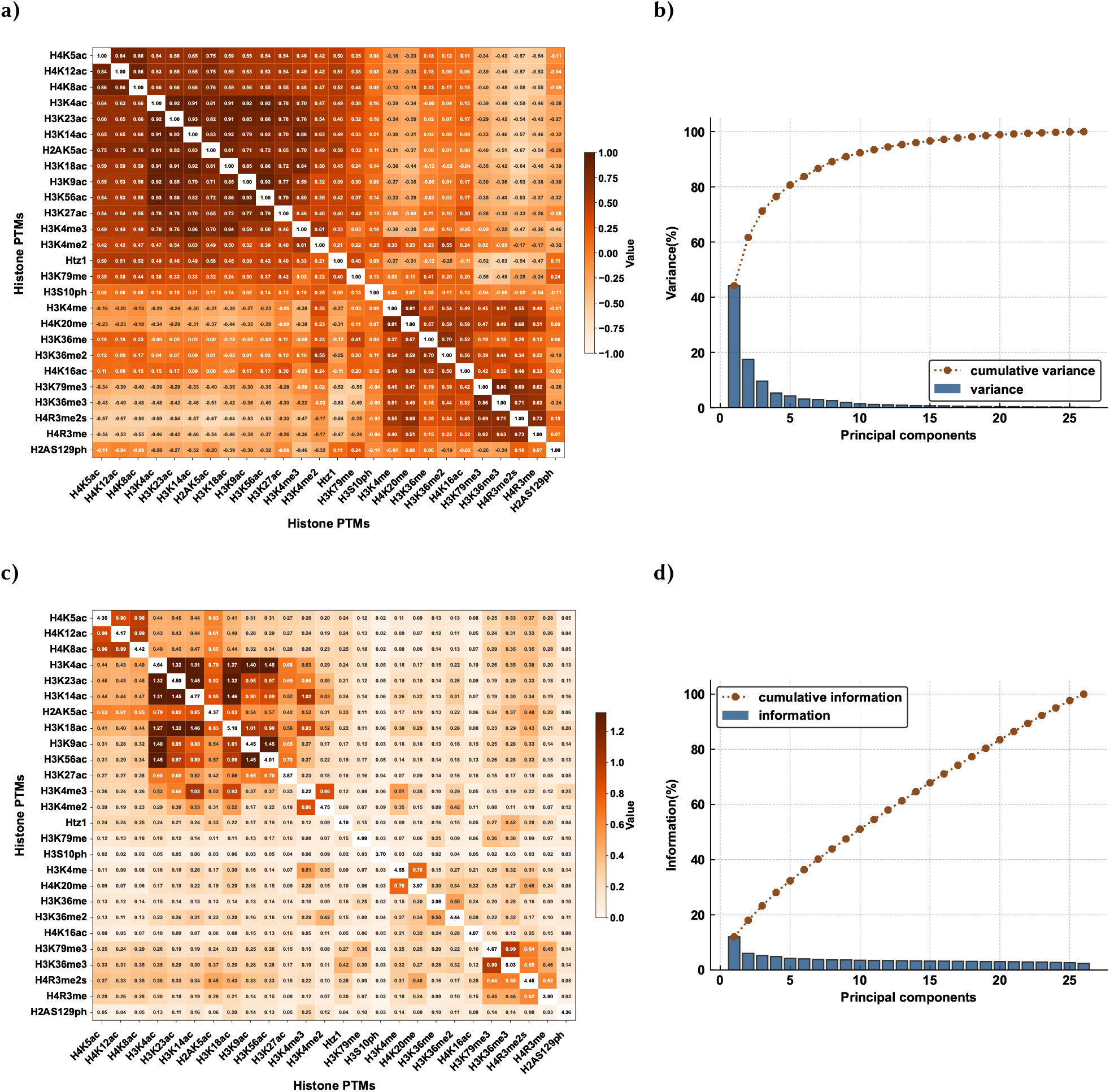
PCA Analysis of Yeast Histone PTMs Dataset: (a) Covariance matrix of the yeast histone PTMs represented on a color scale (light orange = − *ve*, dark orange = + *ve*; diagonal = correlation values), and (b) the corresponding scree plot showing the variance explained by each principal component (bar plot: individual explained variance; line plot: cumulative explained percentage variance). (c) Mutual information (MI) of the pairs of histone PTMs represented as a matrix on a color scale (light orange = low, dark orange = high; diagonal = entropy values), and (d) the corresponding scree plot showing the information explained by each principal component (bar plot: individual explained information; line plot: cumulative explained percentage information).

**Fig. 5.**
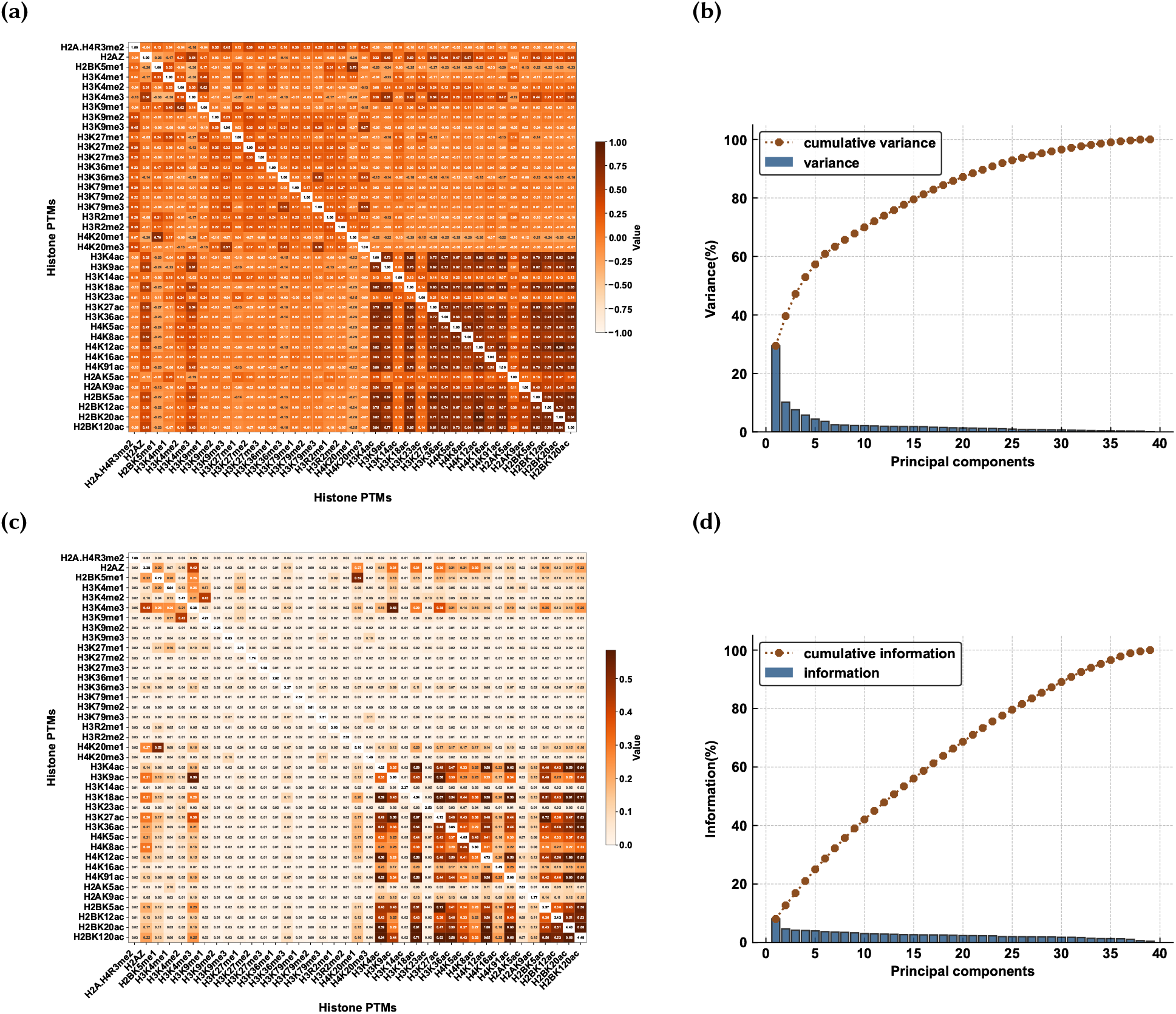
PCA Analysis of Human Histone PTMs Dataset: (a) Covariance matrix of the human histone PTMs represented on a color scale (light orange = −v*e*, dark orange = +*ve*; diagonal = correlation values), and (b) the corresponding scree plot showing the variance explained by each principal component (bar plot: individual explained variance; line plot: cumulative explained percentage variance). (c) Mutual information (MI) of the pairs of histone PTMs represented as a matrix on a color scale (light orange = low, dark orange = high; diagonal = entropy values), and (d) the corresponding scree plot showing the information explained by each principal component (bar plot: individual explained information; line plot: cumulative explained percentage information).

### 3.3. Features identified by MMI help predict histone PTM with high goodness of fit

Extending the pairwise MI further, we computed MMI for triplets of histone PTMs as described in Sec 2.1. Specifically only unique triplets are considered using Eq. 2. For yeast, the MMI values are ranged from −0.49 to 1.2, while for humans, they are ranged from −0.9 to 0.5. The set of all histone PTMs that are part of the lowest MMI triplets (with different thresholds *viz* −0.44, −0.43, −0.41 for the yeast dataset, −0.9,−0.7,−0.6 for the human dataset) are chosen as the principal histones (feature histones). Increasing the threshold will accommodate more feature vectors; these are listed in Table 1. The heatmaps of MMI of the feature histone PTMs in the yeast and human datasets were plotted in Fig. 3(b) and SI Fig. S6(b) respectively.

**Table 1.** Feature histone PTMs identified with different MMI cut-off thresholds.

| Organism | Cut-off | Feature Histones |
| --- | --- | --- |
| Yeast | -0.45 | **H2AS129ph, H3K18ac, H3K36me3, H3K4ac, H4K16ac, H4R3me2s** |
|  | -0.43 | H2AS129ph, H3K18ac, H3K36me3, H3K4ac, H4K16ac, H4R3me2s, **H3K4me, H3K14ac, H3K36me2, H3K4me2, H3K4me3** |
|  | -0.41 | H2AS129ph, H3K18ac, H3K36me3, H3K4ac, H4K16ac, H4R3me2s, H3K4me, H3K14ac, H3K36me2, H3K4me2, H3K4me3, **H4K8ac** |
| Human | -0.90 | **H3K4me1, H3K4me2, H3K4me3** |
|  | -0.70 | H3K4me1, H3K4me2, H3K4me3, **H3K9me1, H4K20me1, H4K91ac** |
|  | -0.60 | H3K4me1, H3K4me2, H3K4me3, H3K9me1, H4K20me1, H4K91ac, **H2BK5me1, H3K18ac, H3K27ac, H4K12ac** |

\* Newly added key histone PCs are highlighted in **bold**.

Using the principal histone PTMs, based on MMI, as features, we predicted the remaining histones with our regression model (Sec. 2.2) employing the *R*^2^ value as a measure of prediction fit (blue-circle line). We also compared the *R*^2^ results obtained from this approach with four other distinct feature selection methods. First, we selected the most highly correlated histone PTM as a feature for each target histone and calculated the square of the Pearson correlation values (red-triangle line), which are equivalent to the *R*^2^ scores in as a univariate linear regression model. For example, in Fig. 6(a) (shown in red), H3K14ac was paired with its most highly correlated histone, H3K23ac (see the correlational heatmap on SI Fig. S7), which had a Pearson correlation coefficent of 0.93 and a squared value of 0.86. As another comparison (purple-diamond line), we combined the feature histone PTMs selected using MMI with the highest correlated histone PTM, H3K23ac (if not originally present in the feature set) and calculated *R*^2^ scores for human dataset. Thirdly (orange-square), we predicted each target PTM using the feature histone PTMs obtained by the OMP feature selection method (for more details, see Sec. 2.3 and SI Sec. *S*8). We also show that the OMP based and pseudo-Inverse based features yielded congruent *R*^2^ coefficients for all the target histone PTMs in SI Figs. 12 (a) and (b). In the final comparison(cyan-x line), for each target histone PTM, all the remaining histone PTMs were used as features and the R^2^ values were computed.

**Fig. 6.**
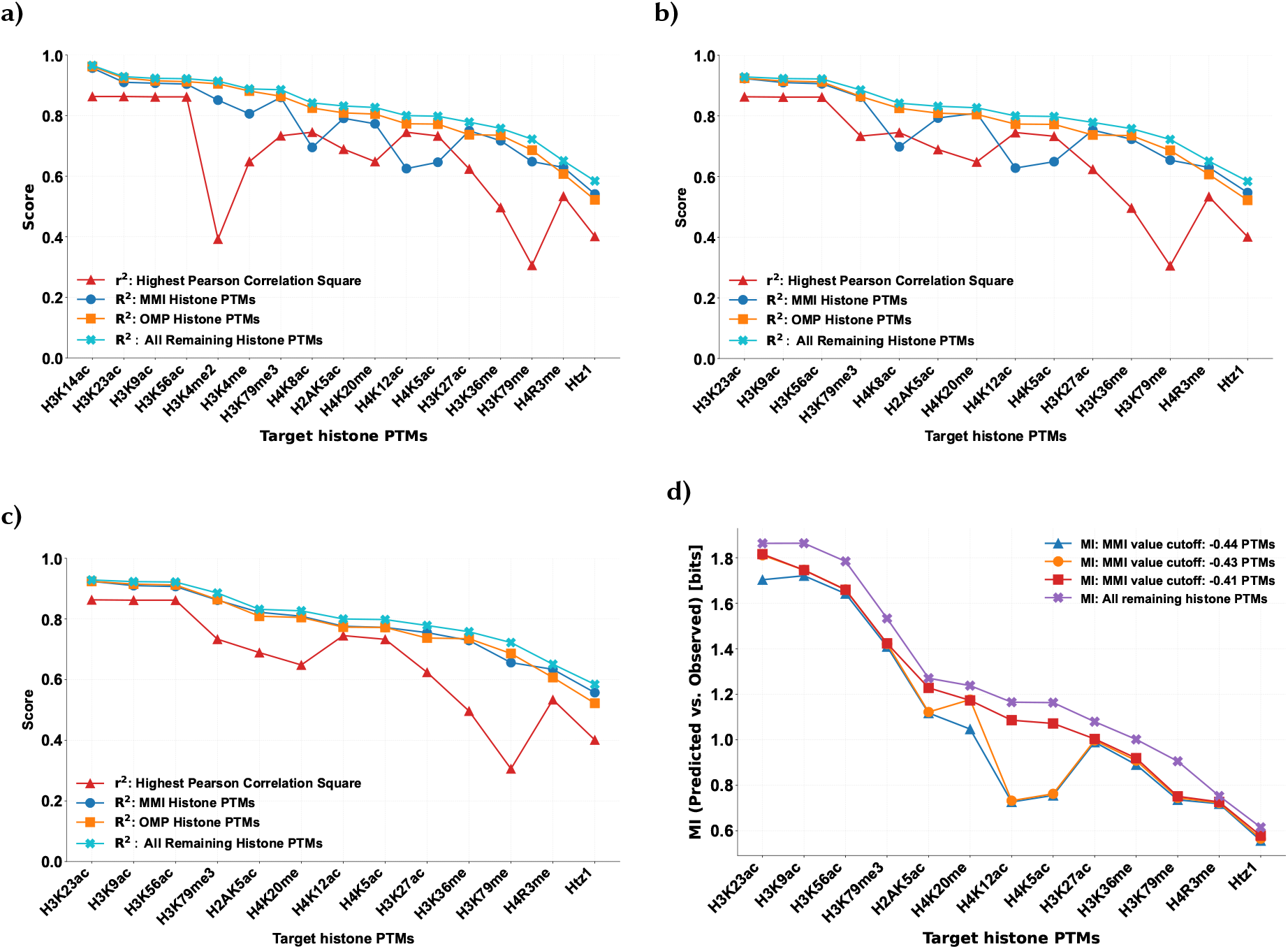
Yeast Histone PTMs - XGBoost Regression Prediction: Plots (a)-(c) illustrate variations in the regression scores calculated under different negative MMI cut-off values (−0.44, −0.43, and −0.41, respectively). Markers: **Red triangles** represent PTMs with the highest squared Pearson correlation values. **Blue circles** indicate PTMs with highly negative MMI values. **orange square** represent the OMP-based feature with R^2^ scores. **cyan X** show the remaining PTMs with the *R*^2^ regression score. (d) illustrates the variations in MI values between predicted and expected target histone PTMs under the above MMI cut-off values.

Figs. 6 and 7 present the regression results for the MMI-based feature selection and its comparison with other feature selection methods from yeast and human datasets respectively. From figs. 6 and 7 ((a), (b), and (c)) it can be observed that as the cut-off values for MMI-based feature selection are relaxed more histone PTMs are moved from the target to the feature set. For the most of the target histone PTMs (except in cases of H3K9me3 and H3K79me3 in human dataset − we expand on these in the SI Sec. S10), the *R*^2^ prediction score using MMI feature selection is higher than those with the highly correlated feature plot. Secondly, when we predicted target histone PTMs using the highly negative MMI histone PTM feature set and highest correlated histone PTMs, the R2 scores closely match those obtained when predicting target histone using all the remaining histone PTMs as features. Similarly, Figs. 6 (d) and 7 (d) show MI values between predicted and original target histone PTMs using the MMI feature sets (at an MMI cut-off of −0.41 for yeast and −0.61 for human), yielding results comparable to those obtained using all remaining histone PTMs.

**Fig. 7.**
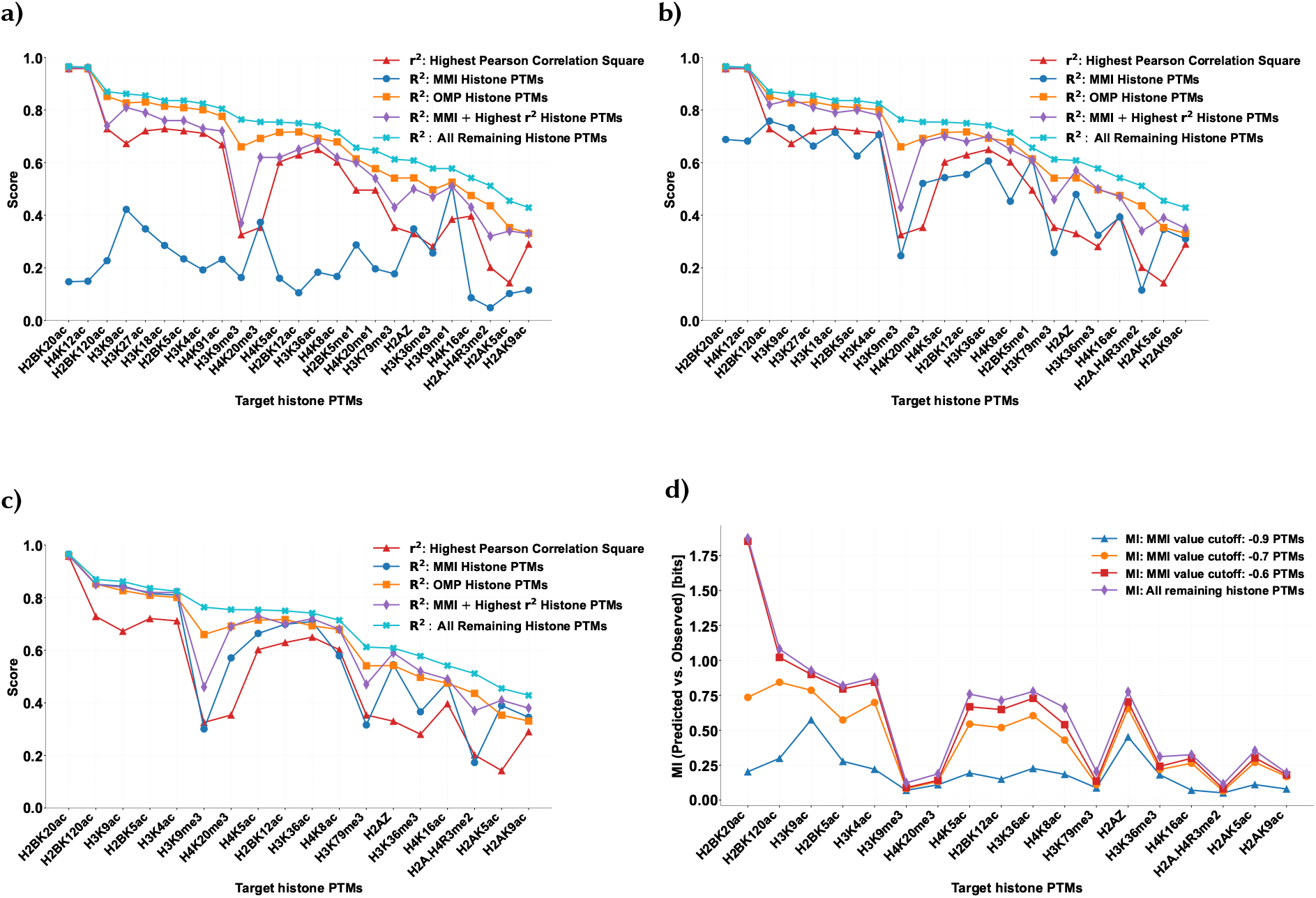
Human Histone PTMs − XGBoost Regression Prediction: Plots (a)-(c) illustrate variations in the regression scores calculated under different negative MMI cut-off values (−0.90, −0.70, and −0.60, respectively). Markers: **Red triangles** represent PTMs with the highest squared Pearson correlation values. **Blue circles** indicate PTMs with highly negative MMI values. **orange square** represent the OMP-based feature with R^2^ scores. **cyan X** show the remaining PTMs with the *R*^2^ regression score. (d) illustrates the variations in MI values between predicted and expected target histone PTMs under the above MMI cut-off values.

## 4. CONCLUSION

Biological datasets are characterized by large dimensionality, diversity and complex relationships between the features (such as genes, proteins, transcription factors, histone PTMs *etc*.) Epigenetic data such as histone PTMs are also set apart by distinct patterns in specific regions of genes and interaction between each other, leading up to the histone code. For prediction of the gene expression based on the patterns in specific histone PTMs across the chromatin, feature selection becomes an important criterion owing to the redundancy and combinatorial complexity among the histone PTMs.

The proposed work introduces a new methodology for feature selection using multivariate mutual information (MMI). We demonstrate this methodology by selecting global set of histone PTMs that can predict the intensities of any other histone PTMs with high R^2^ coefficients. We also compared our method with standard feature selection techniques such as OMP and PCA. It has to be noted that this method is generalizable to any dataset with complex interrelationships between the features.In the future, we intend to extend this method to explore all the other unique relationships in MMI and use them appropriately for explaining the datasets from different biological experiments.

## DATA AND CODE AVAILABILITY

The data analyzed in this study are available in the Gene Expression Omnibus under the accession number GSE61888 and in the Sequence Read Archive under accession number SRA000287. The complete code base is available at https://github.com/nithya-ramakrishnan-biomodeling/multivariate_ptms_analysis-. The corresponding author NR should be contacted for the any other requests.

## ACKNOWLEDGEMENTS

This work was supported by the Department of Electronics, Information Technology, Biotechnology, and Science & Technology of the Government of Karnataka, India.

