## Supplementary material for "Multivariate Mutual Information based Feature Selection for Predicting Histone Post-Translational Modifications in Epigenetic Datasets": SI

### S1 OMP, PCA, and Pseudo Inverse Based Feature Selection Workflow

#### S1.1 OMP-Based Workflow

Orthogonal Matching Pursuit (OMP) is a greedy algorithm; it iteratively selects the most correlated features for each target for feature selection. To compare against MMI-based features, we have applied OMP feature selection methods to identify a subset of features histone PTMs for each target histone PTM. The workflow for feature selection and prediction is illustrated in Fig. S1(a).

#### S1.2 PCA-Based Workflow

As baseline measures, we employed the standard (eigenvector-based) covariance and MI based principal components (PCs) to predict all target histone PTMs. The first 5 PCs obtained from either covariance-based or MI-based approaches (upon suitable transformation) are selected as input features to the XGboost model. The workflow diagram of the selection and prediction process is shown in Fig S1(b).

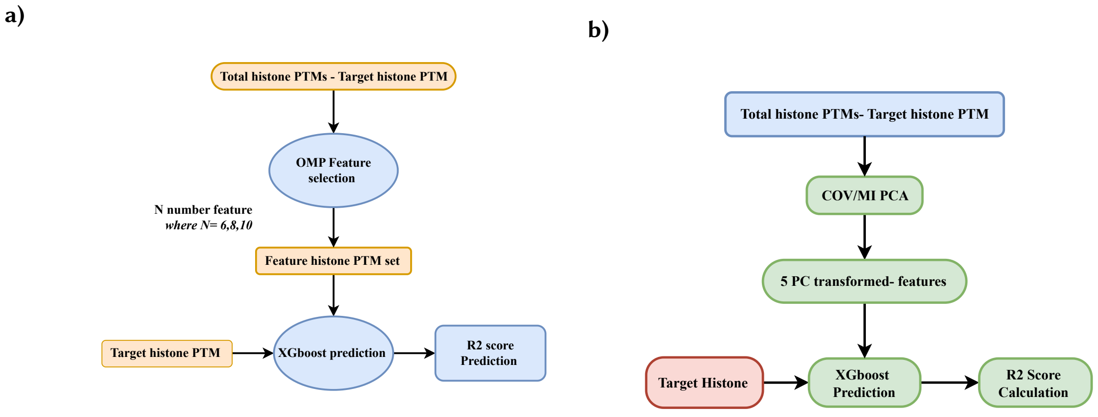


Fig. S1. Workflow of selection of OMP-based and PCA (covariance-based and MI-based) features and prediction of target histones

#### S1.3 Pseudo Inverse Based Workflow

Pseudo Inverse method is employed to obtain the minimum-norm least squares solution for a linear system of the form:

$$\mathbf{y} = A\mathrm{X} \tag{1}$$

In this formulation, $\mathbf{y}$ represents the target histone PTM, $A$ represents the matrix comprising the remaining histone PTMs, and $\mathbf{X}$ is the vector of coefficients indicating the contribution of each histone PTM. $\mathbf{X}$ is estimated by applying the pseudo-inverse of $A$ to $\mathbf{y}$ as follows:

$$\mathrm{X}^{\dagger} = A^{\dagger}\mathbf{y} \tag{2}$$

Following the estimation of $\mathrm{X}^{\dagger}$, feature selection is performed by ranking the histone PTMs based on the absolute values of their corresponding coefficients. In this study, the top 8 histone PTMs with the highest coefficient magnitudes are selected for each target histone PTMs as the selected features.

### S2 Shapiro-Wilk Test on Histone PTM Datasets

To investigate the underlying nature of the data, we applied the Shapiro-Wilk test to each histone PTM distribution in the yeast and human datasets. Tables S1 and S2 present the test **statistic** (second column) and **p-value** (third column) for each histone PTM in the yeast and human datasets respectively. The normality calculation is performed using the *shapiro* function available in the `scipy.stats` library in Python

Table S1. Shapiro-Wilk test results for the yeast histone PTM dataset

| Histone | Statistic | p-value |
|---|---|---|
| H4K5ac | 0.99 | $4.94 \times 10^{-50}$ |
| H4K12ac | 1.00 | $3.95 \times 10^{-27}$ |
| H4K8ac | 1.00 | $5.42 \times 10^{-34}$ |
| H3K4ac | 0.99 | $7.86 \times 10^{-45}$ |
| H3K23ac | 0.99 | $3.33 \times 10^{-46}$ |
| H3K14ac | 0.98 | $6.10 \times 10^{-65}$ |
| H2AK5ac | 0.99 | $8.84 \times 10^{-46}$ |
| H3K18ac | 0.97 | $2.15 \times 10^{-75}$ |
| H3K9ac | 0.99 | $2.06 \times 10^{-50}$ |
| H3K56ac | 0.98 | $2.05 \times 10^{-64}$ |
| H3K27ac | 0.99 | $1.62 \times 10^{-43}$ |
| H3K4me3 | 0.94 | $8.51 \times 10^{-91}$ |
| H3K4me2 | 0.97 | $9.57 \times 10^{-75}$ |
| Htz1 | 0.98 | $1.03 \times 10^{-64}$ |
| H3K79me | 0.99 | $7.48 \times 10^{-56}$ |
| H3S10ph | 0.92 | $2.54 \times 10^{-96}$ |
| H3K4me | 0.99 | $6.68 \times 10^{-59}$ |
| H4K20me | 0.99 | $5.64 \times 10^{-45}$ |
| H3K36me | 0.98 | $7.33 \times 10^{-67}$ |
| H3K36me2 | 0.97 | $4.67 \times 10^{-72}$ |
| H4K16ac | 0.98 | $1.07 \times 10^{-67}$ |
| H3K79me3 | 0.99 | $1.04 \times 10^{-59}$ |
| H3K36me3 | 0.95 | $8.25 \times 10^{-87}$ |
| H4R3me2s | 0.99 | $7.64 \times 10^{-44}$ |
| H4R3me | 1.00 | $3.87 \times 10^{-18}$ |
| H2AS129ph | 0.89 | $1.06 \times 10^{-105}$ |

Table S2. Shapiro-Wilk test results for the human histone PTM dataset

| Histone Modification | Statistic | p-value |
|---|---|---|
| H2A.H4R3me2 | 0.46 | $1.19 \times 10^{-179}$ |
| H2AZ | 0.59 | $5.16 \times 10^{-170}$ |
| H2BK5me1 | 0.73 | $1.18 \times 10^{-155}$ |
| H3K4me1 | 0.84 | $2.49 \times 10^{-140}$ |
| H3K4me2 | 0.85 | $1.35 \times 10^{-138}$ |
| H3K4me3 | 0.74 | $5.52 \times 10^{-155}$ |
| H3K9me1 | 0.91 | $1.88 \times 10^{-122}$ |
| H3K9me2 | 0.57 | $1.42 \times 10^{-171}$ |
| H3K9me3 | 0.14 | $1.36 \times 10^{-196}$ |
| H3K27me1 | 0.88 | $8.39 \times 10^{-131}$ |
| H3K27me2 | 0.48 | $3.51 \times 10^{-178}$ |
| H3K27me3 | 0.46 | $1.05 \times 10^{-179}$ |
| H3K36me1 | 0.80 | $5.17 \times 10^{-146}$ |
| H3K36me3 | 0.45 | $2.08 \times 10^{-180}$ |
| H3K79me1 | 0.70 | $3.08 \times 10^{-160}$ |
| H3K79me2 | 0.33 | $1.83 \times 10^{-187}$ |
| H3K79me3 | 0.41 | $1.47 \times 10^{-182}$ |
| H3R2me1 | 0.78 | $3.45 \times 10^{-150}$ |
| H3R2me2 | 0.58 | $9.11 \times 10^{-171}$ |
| H4K20me1 | 0.74 | $4.67 \times 10^{-155}$ |
| H4K20me3 | 0.19 | $4.54 \times 10^{-194}$ |
| H3K4ac | 0.65 | $4.23 \times 10^{-165}$ |
| H3K9ac | 0.55 | $9.92 \times 10^{-174}$ |
| H3K14ac | 0.77 | $1.81 \times 10^{-150}$ |
| H3K18ac | 0.71 | $4.97 \times 10^{-159}$ |
| H3K23ac | 0.78 | $3.57 \times 10^{-149}$ |
| H3K27ac | 0.69 | $4.13 \times 10^{-161}$ |
| H3K36ac | 0.57 | $4.11 \times 10^{-172}$ |
| H4K5ac | 0.69 | $3.43 \times 10^{-161}$ |
| H4K8ac | 0.68 | $1.09 \times 10^{-161}$ |
| H4K12ac | 0.73 | $9.75 \times 10^{-157}$ |
| H4K16ac | 0.73 | $1.45 \times 10^{-156}$ |
| H4K91ac | 0.72 | $3.48 \times 10^{-157}$ |
| H2AK5ac | 0.72 | $5.35 \times 10^{-157}$ |
| H2AK9ac | 0.51 | $1.15 \times 10^{-176}$ |
| H2BK5ac | 0.55 | $3.18 \times 10^{-173}$ |
| H2BK12ac | 0.62 | $1.78 \times 10^{-167}$ |
| H2BK20ac | 0.69 | $3.28 \times 10^{-161}$ |
| H2BK120ac | 0.65 | $1.26 \times 10^{-164}$ |

### S3 Histone PTM Distribution

The distribution of the yeast and human PTM datasets before and after the zero-mean normalization are shown in figures S2, S3, S4, and S5, respectively. For the human dataset, extreme values(greater than 40) were clipped.

#### S3.1 Yeast Raw Data Distribution

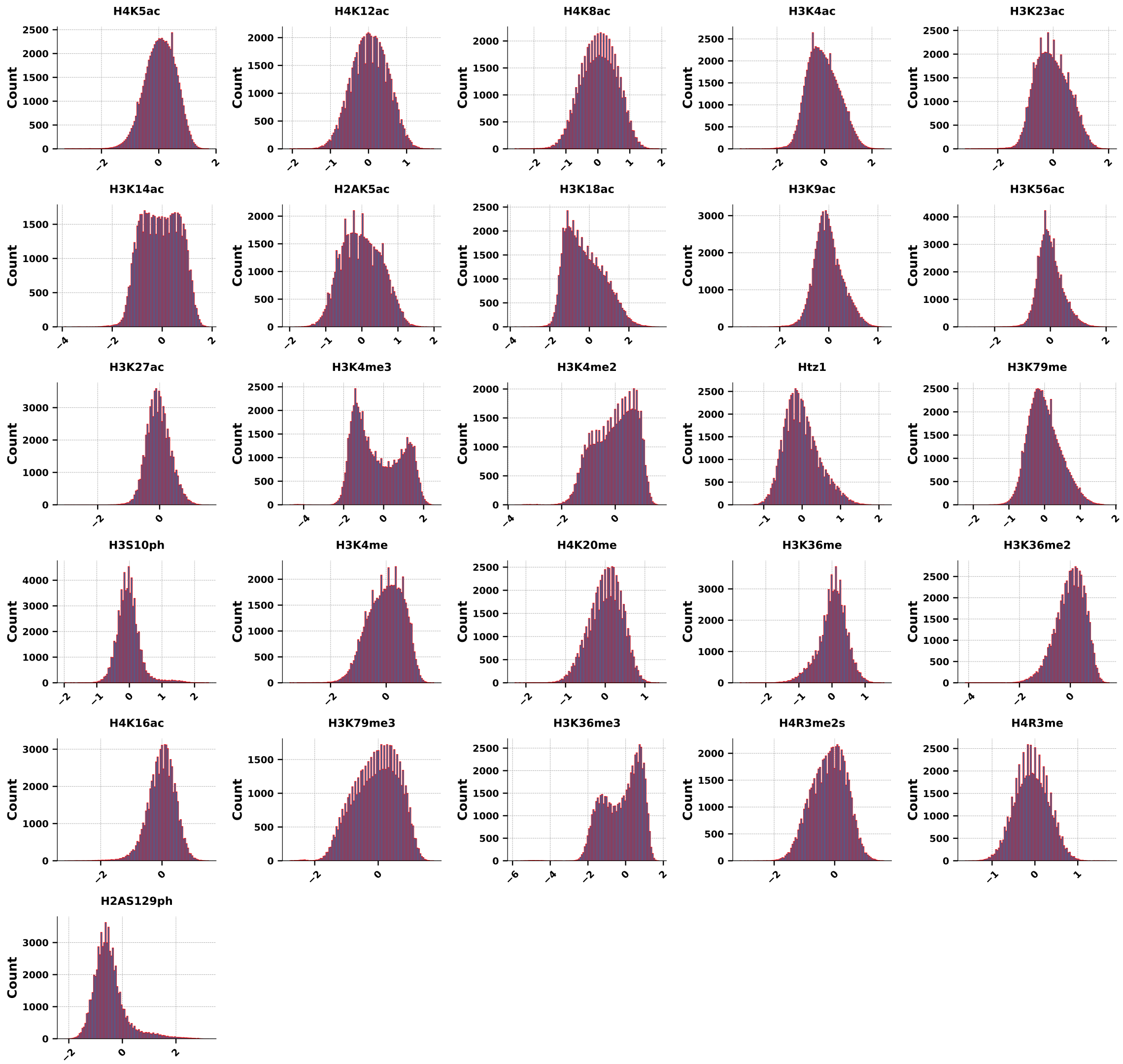


Fig. S2. Histogram distributions of yeast histone PTM dataset before zero-mean normalization

### S3.2 Yeast Data Zero-Mean Distribution

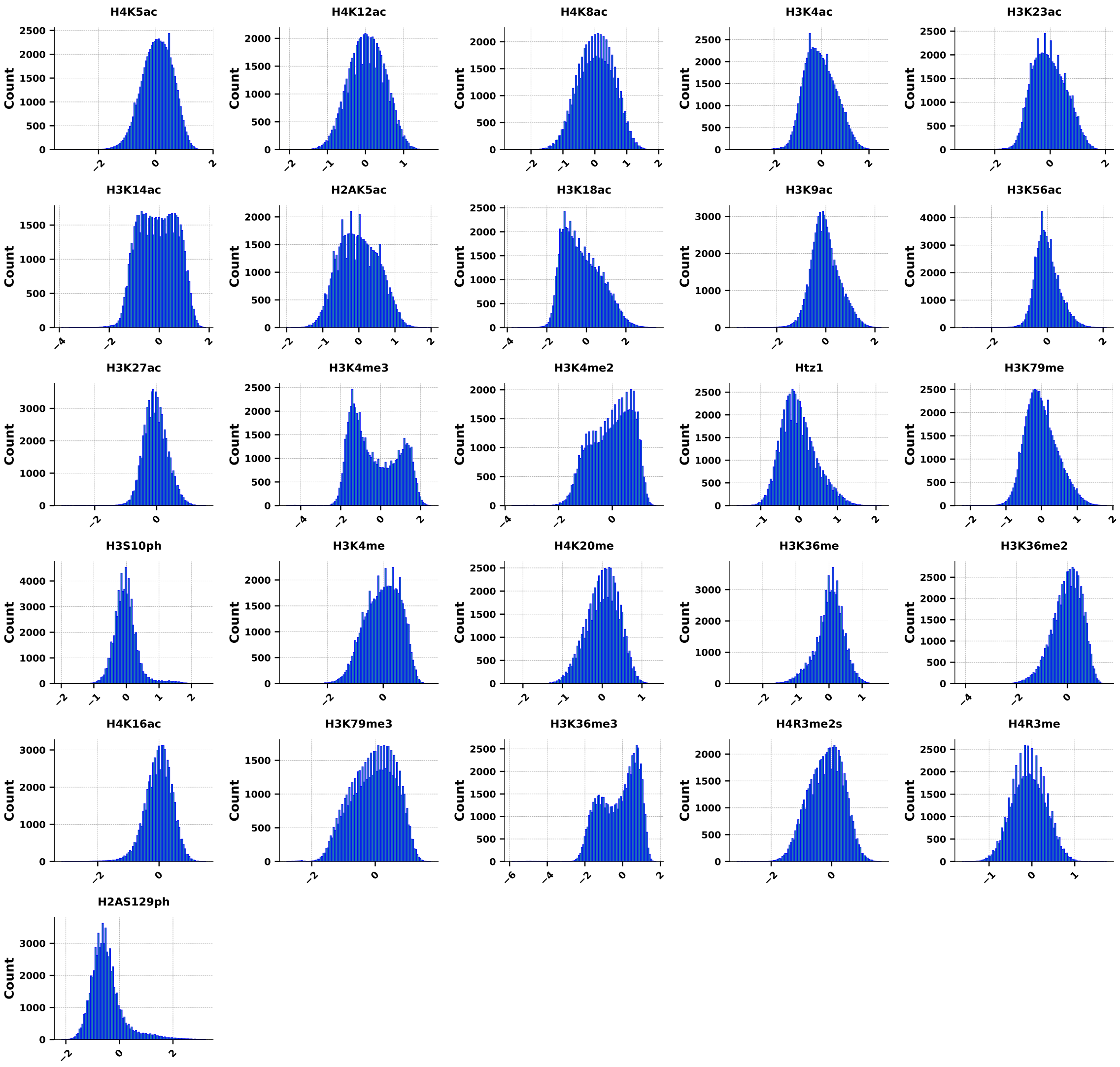


Fig. S3. Yeast histone PTMs zero-mean data

### S3.3 Human Raw Data Distribution

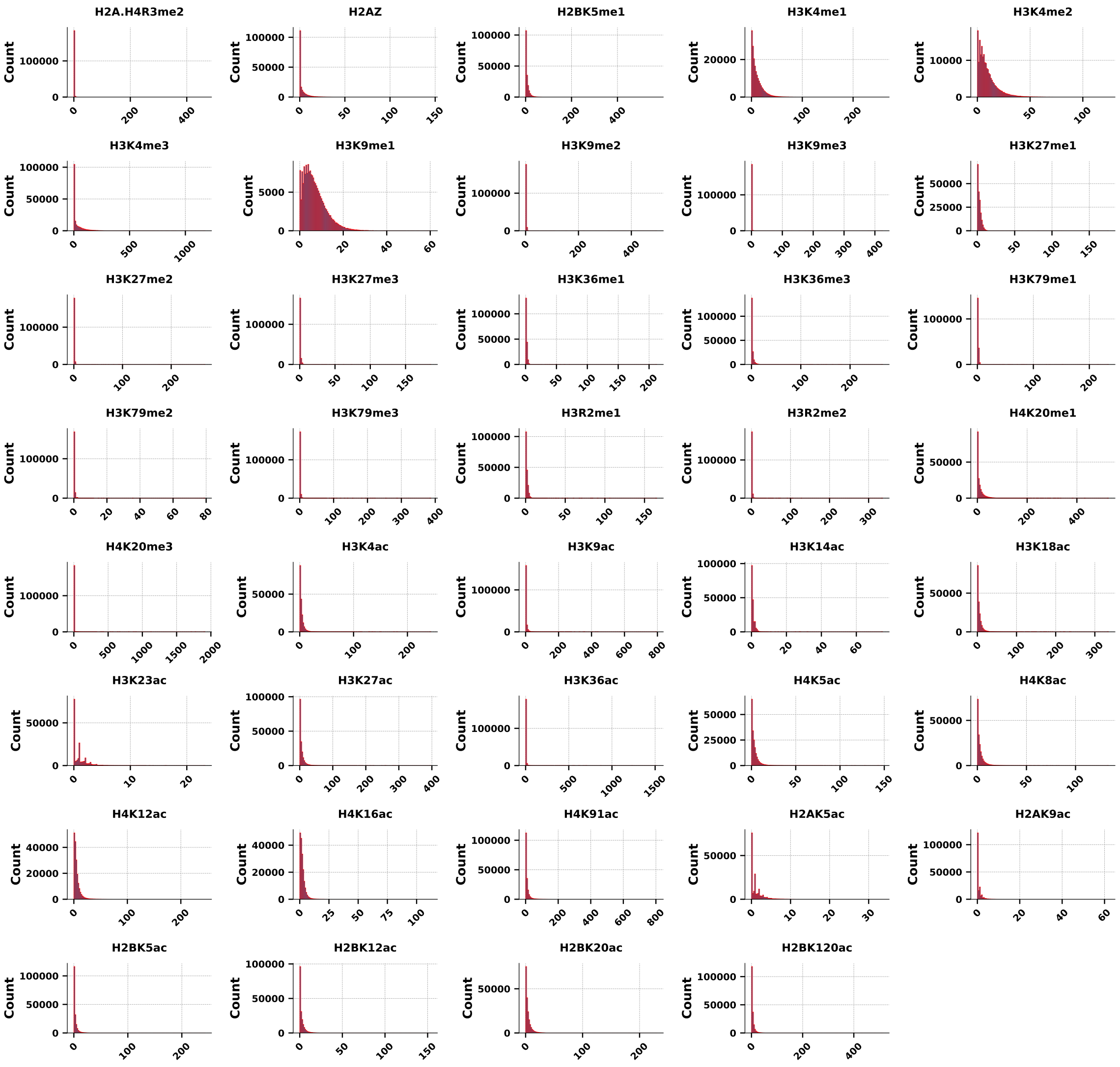


Fig. S4. Human histone PTMs raw data

### S3.4 Human Zero-Mean Distribution

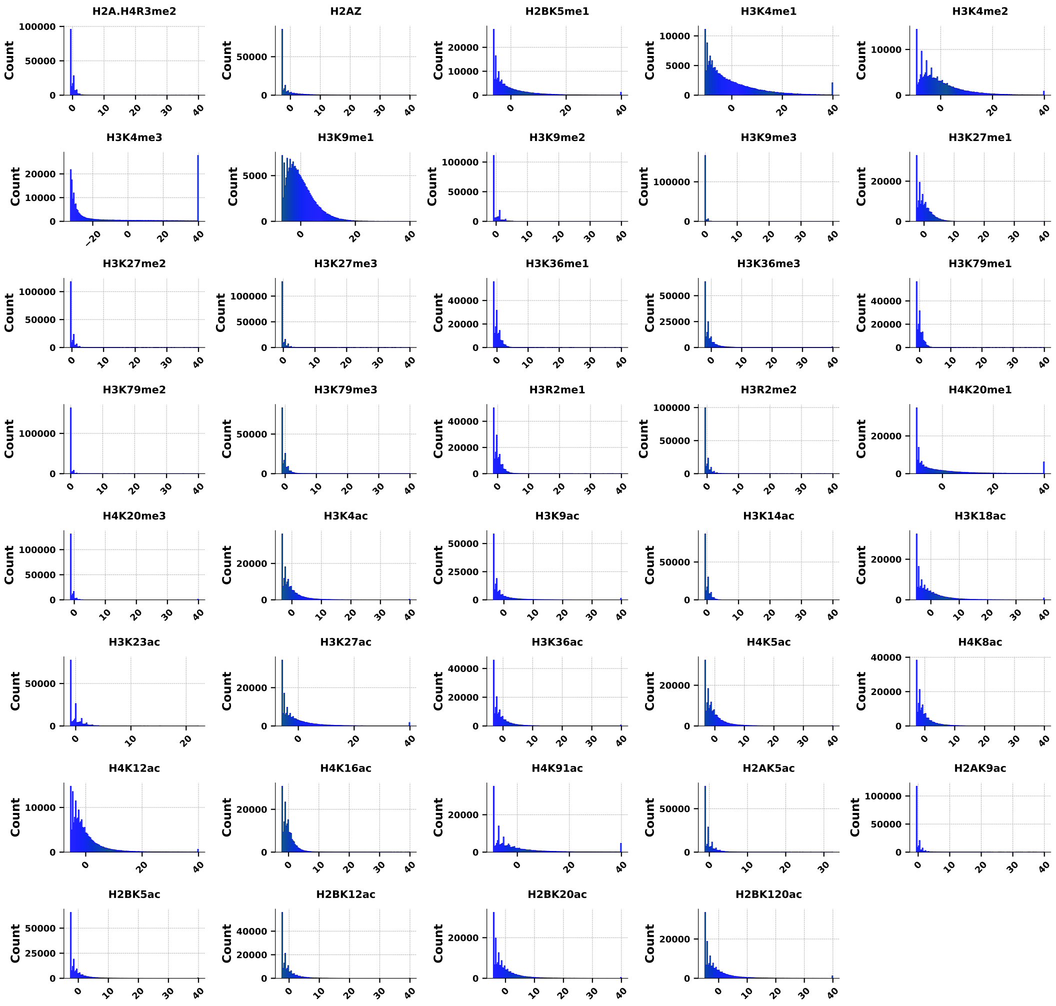


Fig. S5. Human histone PTMs zero-mean data

### S4 Variations of Entropy with Different Binning Size in Yeast and Human Datasets

The entropy of each histone PTM in both datasets was computed based on binned probability distributions. To determine the optimal bin width for entropy estimation, we evaluated entropy values across a range of bin sizes and selected the most suitable width for subsequent analysis on both datasets. Both the figs. S6 (a) and (b) shows entropy variations for the yeast and human datasets respectively.

**(a)**

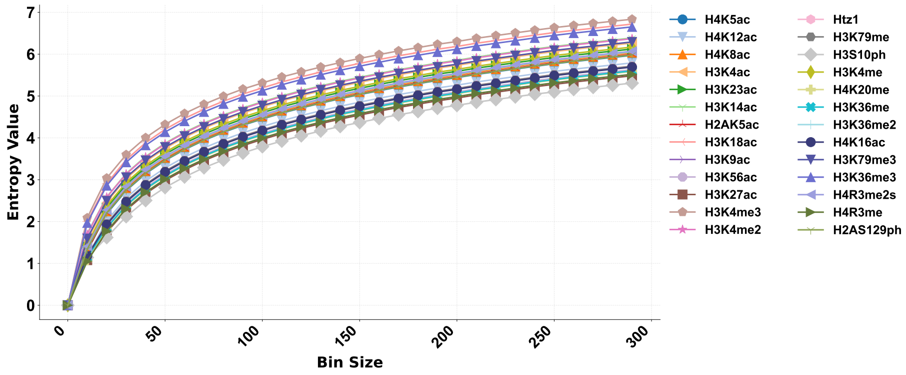


**(b)**

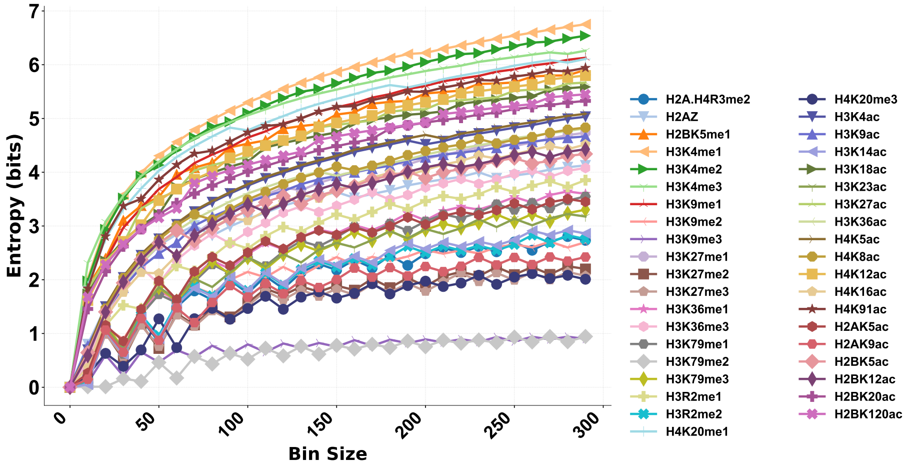


Fig. S6. **Entropy value of the histone PTMs using different bin sizes:** The lineplots (a) and (b) show entropy values of the each histone PTM in the yeast and human datasets respectively.

### S5 Prediction of Histone PTMs Based on PCA-Based Features

The yeast and human PTMs values are predicted using XGboost models with the PCA-based (both covariance and MI based) features as input distributions. The resulting $R^2$ coefficients are shown in figures respectively. For comparison, we also obtained the results with the highest correlated features and remaining features (each target histone PTM).

**(a)**

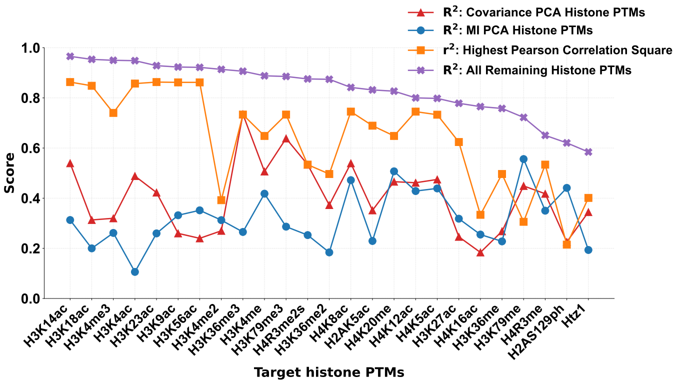


**(b)**

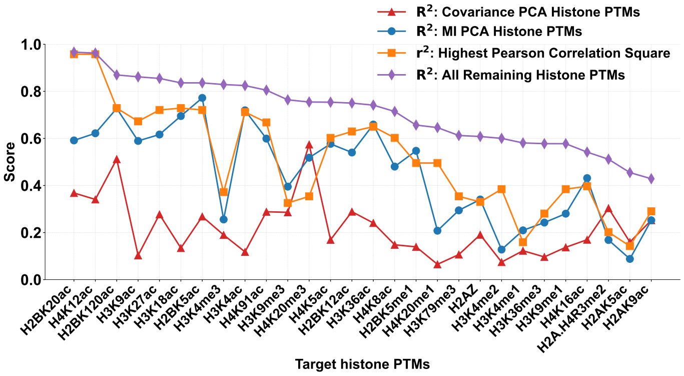


Fig. S7. $R^2$ **values of the XGboost prediction models using PCA features:** The lineplots (a) and (b) show the comparison results of yeast and human PCA based features predictions respectively.

### S6 Feature Selection Using OMP and MMI in Human Dataset

OMP based feature selection method was applied to each target histone PTM to identify correlated features in the human dataset, as shown in Fig. S8(a). The x-axis represents the target histone PTMs, while the y-axis denotes the selected feature histone PTMs for each target. The color scheme indicates the presence (red) or absence (white) of a feature histone PTM.

Similarly, negative MMI histone triplets are illustrated in Fig. S8 (b), where the x-axis corresponds to the first and second histone PTMs, and the y-axis indicates the third PTM in each triplet. The MMI values are encoded using the colour map shown beside the figure.

**a)**

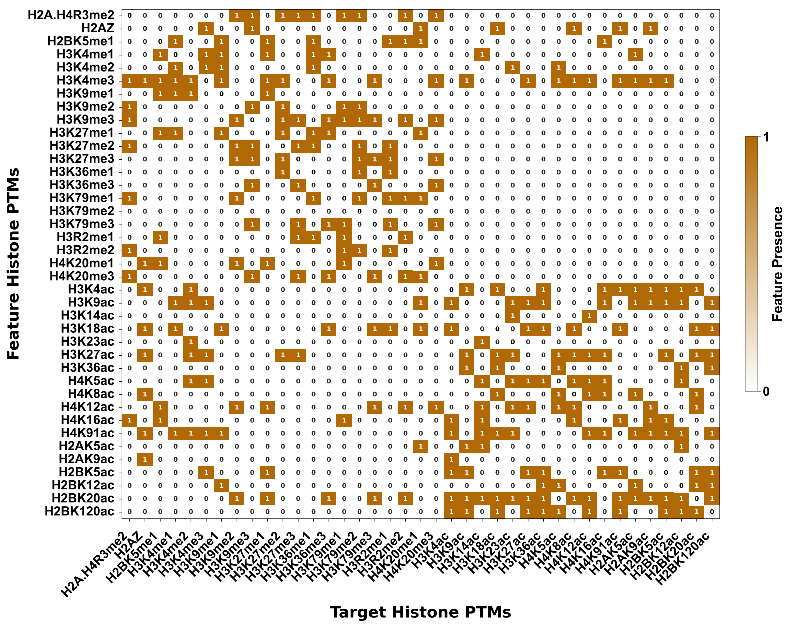


**b)**

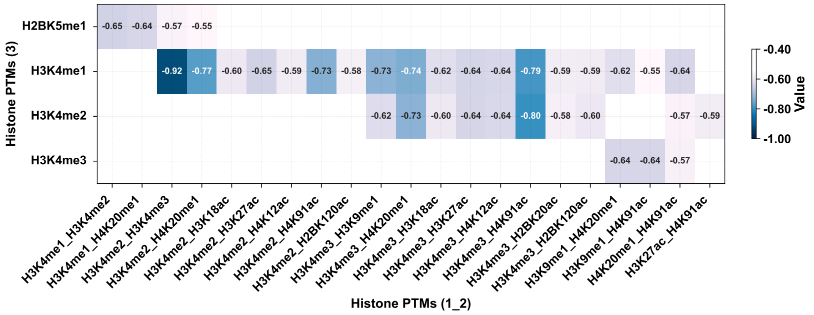


Fig. S8. **Feature Selection using OMP and MMI in human dataset**: Plots (a) and (b) represent the features that are obtained from the OMP and MMI techniques on human dataset respectively.

### S7 Pearson Correlational Results between Histone PTMs

The Pearson correlational heatmaps between the yeast and human histone PTMs are shown in Figs. S9 and S10, respectively.

#### S7.1 Yeast Correlational Heatmap

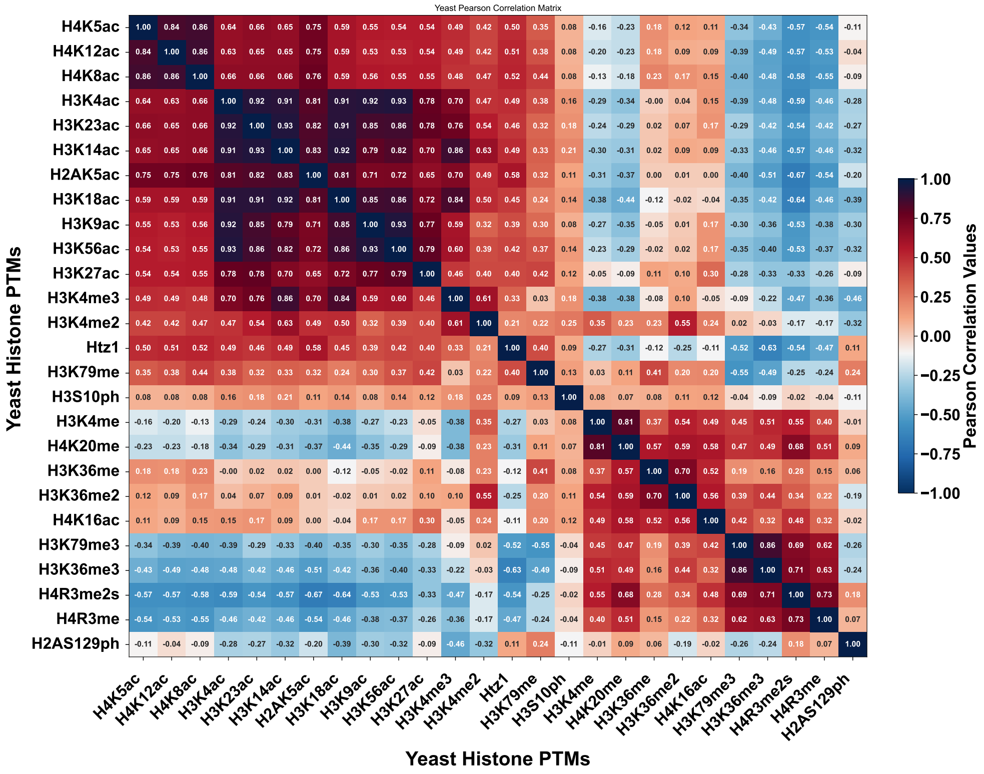


Fig. S9. Correlation heatmap of yeast dataset

### S7.2 Human Correlational Heatmap

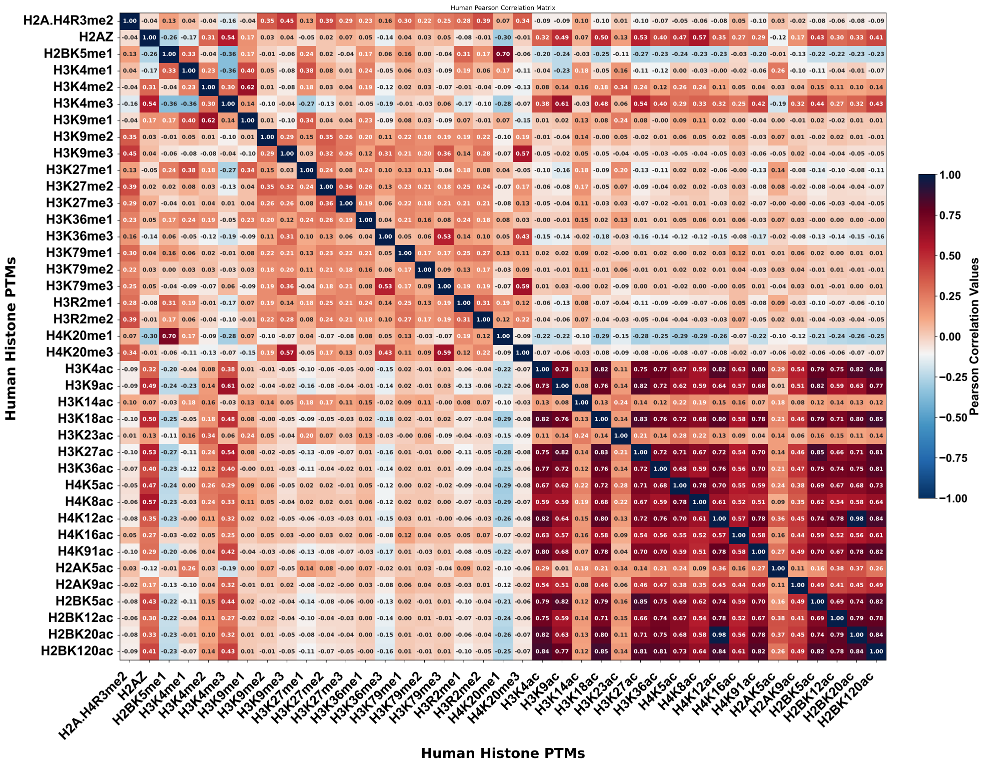


Fig. S10. Correlation heatmap of human dataset

### S8 Goodness of Fit Comparison between the OMP-Based Target Histone Prediction

For the OMP-based target histone target predictions, we chose three different cardinalities feature sets ($n = 6, 8, 10$). The feature selection work flow is described in SI Sec.$S1$ The results were then compared with two other different prediction score methods that use all remaining histones as features and the highest correlated histone squared. The comparison results are depicted in Fig. S11.

**(a)**

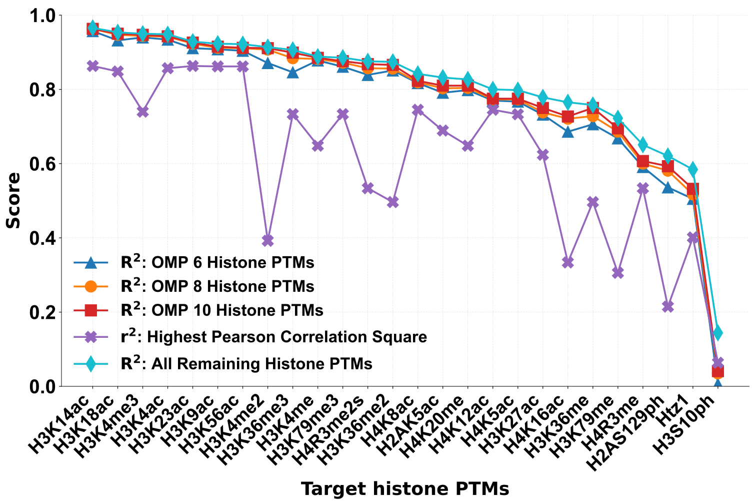


**(b)**

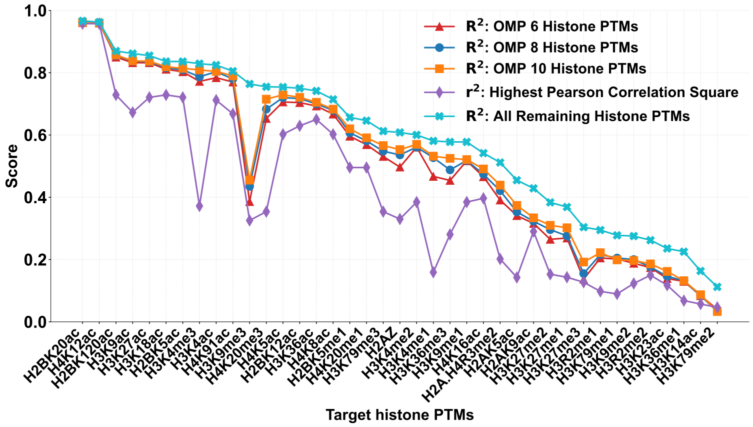


Fig. S11. **$R^2$ score comparison with OMP features:** The lineplots (a) and (b) show the comparison results of yeast and human OMP based features predictions respectively.

The $R^2$ prediction scores using 6, 8, and 10 features selected by the OMP method are represented by blue, orange, and red bars, respectively. The $R^2$ prediction scores based on the high correlation and all remaining histones are represented by purple and cyan bars.

### S9 Goodness of Fit Comparison between OMP and Pseudo Inverse Based Feature Histone PTMs

The R2 coefficients for prediction each target histone PTMs, using feature histone PTMs selected through OMP and pseudo-inverse techniques, are plotted in Figs. S12 (a) and (b) for the yeast and human datasets, respectively.

**(a)**

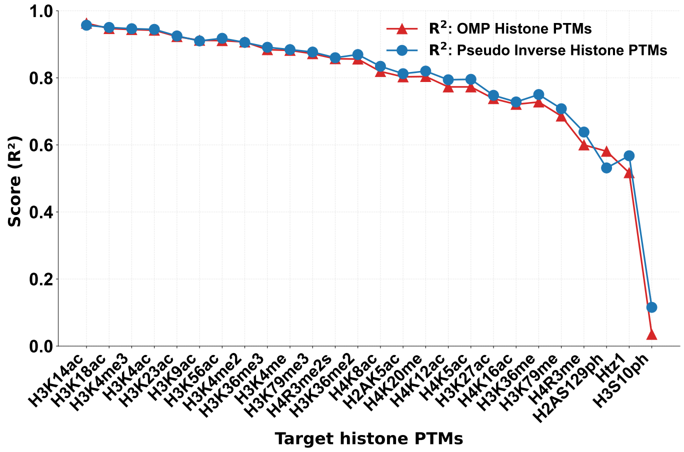


**(b)**

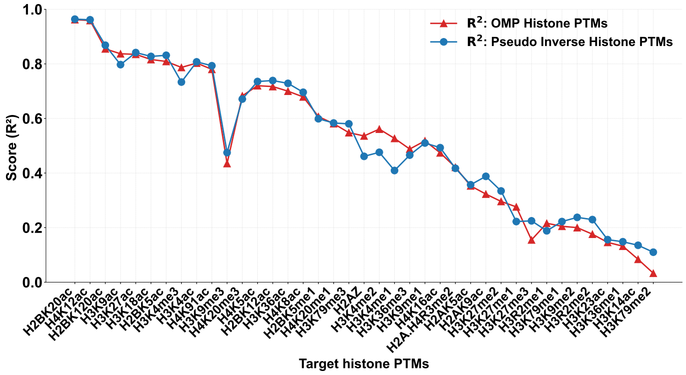


Fig. S12. $R^2$ **scores comparison of the regression results between OMP and Pseudo Inverse based selected feature** The lineplots (a) and (b) show $R^2$ scores comparison results of histone PTMs in the yeast and human datasets respectively.

### S10 Regression Prediction of Human Histone PTMs Outliers: H3K9me3 and H3K79me3

To analyze the behaviour of the outliers in the human dataset, we predicted H3K9me3 and H3K79me3 using each of the remaining histone PTMs and plotted the $R^2$ values individually and cumulatively along with MMI histone feature histone PTMs at $-0.6$ MMI value cut-off. It can be observed that the highest $R^2$ value for H3K9me3 was with H2AZ and H3K36me3 for H3K79me3. One could note that H2AZ and H3K36me3 are not part of the MMI-based feature-set.

**(a)**

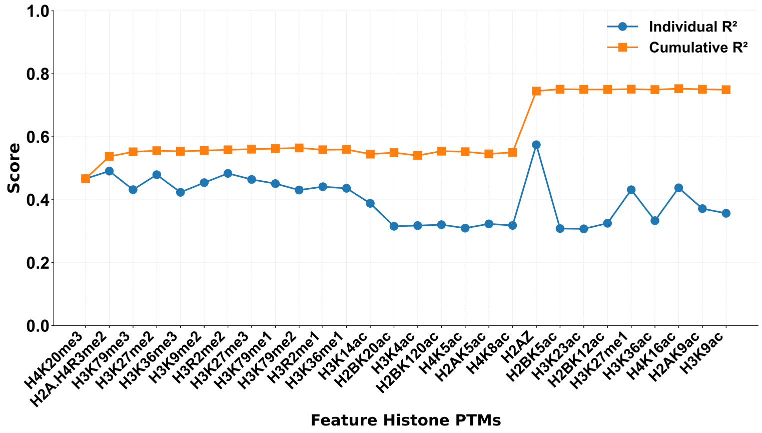


**(b)**

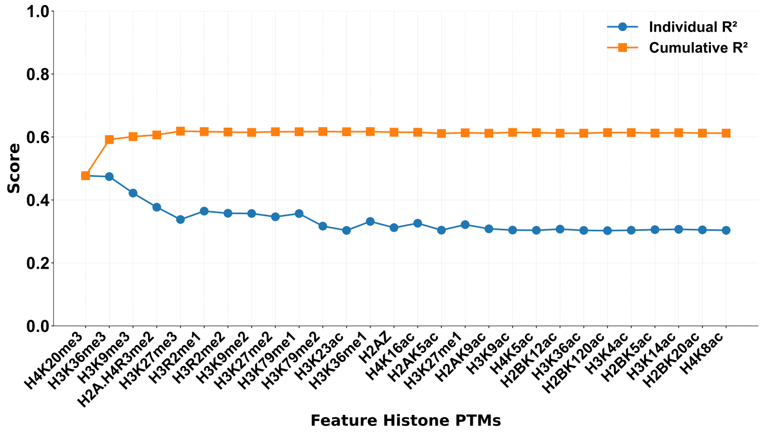


Fig. S13. **Regression based prediction of human histone PTMs (a) H3K9me3 and (b) H3K79me3 using remaining histones as features** Feature histones on the x-axis is ordered from highest to lowest correlated to the target histone PTM. The **red line** indicate that $R^2$ score of the cumulative prediction while the the **blue line** shows the $R^2$ of the individual feature prediction
